# A negative feedback loop between small GTPase Rap1 and mammalian tumor suppressor homolog KrsB regulates cell-substrate adhesion in *Dictyostelium*

**DOI:** 10.1101/2024.11.07.622562

**Authors:** Yulia Artemenko, Gengle Niu, Megan E. Arnold, Kelsey E. Roberts, Bianca N. Fernandez, Tiffany Flores, Harper D. McClave, Michael Paestella, Jane Borleis, Peter N. Devreotes

## Abstract

Cell adhesion to the substrate influences a variety of cell behaviors and its proper regulation is essential for migration, although details of the molecular pathways regulating cell adhesion during migration are lacking. Rap1 is a small GTPase that regulates adhesion in mammalian cells, as well as in *Dictyostelium discoideum* social amoeba, which is an established model for studying directed cell migration. In *Dictyostelium*, Rap1 controls adhesion via its effects on adhesion mediator talin and Ser/Thr kinase Phg2, which inhibits myosin II function. Kinase responsive to stress B (KrsB), a homolog of mammalian tumor suppressor MST1/2 and *Drosophila* Hippo, also regulates cell adhesion and migration, although the molecular mechanism of KrsB action is not understood. Since KrsB has been shown to interact with active Rap1 by mass spectroscopy, we investigated the genetic interaction between Rap1 and KrsB. Cells lacking KrsB have increased adhesion to the substrate, which leads to reduced movement. Expression of constitutively active Rap1 G12V increased cell spreading and adhesion even in the absence of KrsB, suggesting that Rap1 does not require KrsB to mediate cell adhesion. In contrast, dominant negative Rap1 S17N completely reversed the over-adhesive phenotype of KrsB-null cells and impaired KrsB phosphorylation, suggesting that KrsB activation and function in adhesion requires Rap1. Even though Rap1 did not require KrsB for its function in adhesion, KrsB negatively regulates Rap1 function as seen by increased cortical localization of Rap1 in KrsB-null cells. Consistent with this, chemoattractant-induced activation of downstream effectors of Rap1, TalB and Phg2, was increased in the absence of KrsB. Taken together, these findings suggest that Rap1 leads to activation of KrsB, which inhibits Rap1 and its downstream targets, shutting off adhesion. The existence of a negative feedback loop between Rap1 and KrsB may contribute to the dynamic regulation of cell adhesion that is necessary for rapid amoeboid-type migration.

## Introduction

Proper cell-substrate adhesion is critical for migration since it allows a cell to generate traction that is required for forward movement. Cells undergoing mesenchymal-type migration, such as fibroblasts, rely on focal adhesions, which mediate cell attachment to the extracellular matrix and are formed by clusters of integrins interacting with a number of intracellular components, including vinculin, talin, paxillin, focal adhesion kinase, and actin cytoskeleton (Parlani et al., 2023; Pourjafar and Tiwari, 2024). In contrast, cells undergoing amoeboid-type migration, such as leukocytes or metastatic cancer cells, rely on weak adhesion to substrate, allowing the cells to migrate very quickly. These cells form transient short-lived integrin complexes and can even migrate in the absence of integrins, especially in three-dimensional environments (Smith et al., 2005; Lämmermann et al., 2008). Although we know many components of the signal transduction network that regulates migration, details of the pathways regulating adhesion, especially in the absence of integrins, are lacking.

*Dictyostelium discoideum* social amoeba has been instrumental for our understanding of the signal transduction network driving directed migration, with many of the pathways conserved in mammalian cells undergoing amoeboid-type movement (Artemenko et al., 2014). *D. discoideum* spends most of its life within the growth (vegetative) phase, as haploid amoeba, if provided with enough nutrition (Bozzaro, 2019). During this stage cells depend on chemotaxis toward signals released from its food sources (bacteria or yeast), such folic acid. When food is scarce, cells enter the differentiation phase and aggregate by chemotaxing toward a source of cyclic AMP (cAMP) to form a series of multicellular structures leading to the formation of a fruiting body containing spores.

Although *D. discoideum* cells possess a protein that is similar to beta-integrin, SibA, as well as homologs of talin and paxillin, they do not form typical focal adhesions (Bukharova et al., 2005; Cornillon et al., 2008; Tsujioka et al., 2008). Instead, *D. discoideum* cells rely on non-specific van der Waals interactions for binding to various substrates, as well as form transient attachment sites similarly to leukocytes (Loomis et al., 2012; Mijanović and Weber, 2022). In fact, both paxillin and talin localize to distinct puncta on the basal surface of *D. discoideum* cells (Hibi et al., 2004; Bukharova et al., 2005; Tsujioka et al., 2008). Thus, *D. discoideum* offers a unique system to study integrin-independent migration. Exploring the mechanisms of how *D*. *discoideum* migrates and adheres to substrate could provide key insight into how other amoeboid cells migrate.

Small GTPase Rap1 is one of the few well-established regulators of adhesion and chemotaxis both in *D. discoideum* and mammalian cells (Hilbi and Kortholt, 2019). In *D. discoideum* chemoattractant binding to its G protein-coupled receptor leads to activation of the Rap1 pathway. Ser/Thr kinase Phg2, which leads to Myosin II disassembly at the leading edge, and cytoskeletal adapter protein Talin B, which is important for adhesion, are downstream targets that are regulated by Rap1 (Jeon et al., 2007; Plak et al., 2016). Activation of Rap1 also affects actin polymerization and pseudopod extension, allowing migration of the cells. Overexpression of Rap1 results in cytoskeletal defects and flattened cell morphology (Jeon et al., 2007).

Another regulator of adhesion in *D. discoideum* is KrsB (Kinase Responsive to Stress B), which is a homolog of mammalian tumor suppressor MST1/2 and *Drosophila* Hippo (Artemenko et al., 2012). KrsB is activated downstream of the chemoattractant cAMP by autophosphorylation with similar dynamics as activation of other leading-edge proteins. KrsB is a negative regulator of cell spreading and substrate attachment, and KrsB-null cells showed increased adhesion and reduced directional movement; however, the molecular mechanism of KrsB action is not understood. Several lines of evidence suggest that KrsB may interact with the pathway involving Rap1 or its effectors, including an interaction between KrsB and active Rap1 that was detected by mass spectrometry, as well as phenotypic similarity between cells lacking KrsB and those expressing constitutively active Rap1 or its activator, GbpD (Jeon et al., 2007; Artemenko et al., 2012; Plak et al., 2016). Additionally, mammalian homolog of KrsB, Mst1, has been shown to alter T lymphocyte adhesion and polarization in a Rap1-dependent manner (Katagiri et al., 2006). In this study we investigated the genetic interaction between Rap1 and KrsB and found that KrsB acts as a negative regulator of the Rap1 pathway.

## Materials and Methods

### Cell Culture

*D. discoideum* cells were cultured on plates or in shaking culture in HL-5 media at 20°C according to standard conditions (Artemenko et al., 2011). Wild-type (Ax2) cells were generously provided by R. Kay (MRC Laboratory of Molecular Biology, Cambridge, United Kingdom). Cells lacking KrsB (*krsB^-^*) were previously generated in (Artemenko et al., 2012) and stocks were routinely maintained in the presence of 10 μg/mL blasticidin S sulfate to ensure lack of cross-contamination with wild-type cells. Transformation of cells was performed with H-50 Buffer using a Gene Pulser Xcell electroporator (Bio-Rad) according to standard procedures (Gaudet et al., 2007). Transformed cell lines were selected using 20 µg/ml G418 and/or 50 µg/ml hygromycin for at least one week. Success of the transformation was verified by imaging cells in developmental buffer (DB; phosphate buffer supplemented with 2 mM MgSO_4_ and 0.2 mM CaCl_2_) under 630X magnification with brightfield illumination and epifluorescence with a GFP or an RFP filter set using a Zeiss LSM700 confocal microscope, and/or immunoblotting cell lysates with appropriate antibodies (see below).

Myc-tagged Rap1 constructs in the pEXP4(+) vector were generously provided by R. Firtel (University of California San Diego). Rap1 constructs tagged with mRFPmars in the pDM318 vector were generated in this study. pDM318 plasmid carrying an RFP tag and GFP-tagged RalGDS in pDM115 plasmid were generously provided by A. Kortholt (University of Groningen). GFP-tagged TalB and TalA, Phg2 in pDXA-GFP vector (pFL712), as well as pREP, which is needed for co-transformation with pDXA-GFP, were obtained from the Dicty Stock Center (Chicago, IL, United States).

### Cloning of RFP-tagged Rap1 constructs

Full-length Rap1 or Rap1 G12V was subcloned from the pEXP4(+)/Rap1 or pEXP4(+)/Rap1 G12V plasmid into the pDM318 vector, which includes an N-terminal RFP tag, using standard molecular cloning methods. Rap1 or Rap1 G12V without the initial ATG sequence was amplified by PCR with Phusion DNA polymerase (Thermo Fisher Scientific) using primers with BamHI and SpeI restriction sites (5’ CCCGGATCCCCTCTTAGAGAATTCAAAATCGTC 3’ and 5’ CGGACTAGTTTACAATAAAGCACATTTTGATTTAGC 3’). The insert was cut with BamHI and SpeI and ligated into pDM318 vector pre-cut with <u>BglII</u> and <u>SpeI</u> using T4 DNA ligase (Thermo Fisher Scientific). Ligation products were transformed into NEB 5-alpha competent *E. coli*. Transformed cells were plated on LB plates with 150 µg/ml ampicillin and incubated at 37°C for at least 16 hours. Plasmid extraction was performed using GeneJET^TM^ Plasmid Miniprep Kit (Thermo Fisher Scientific). Colonies were screened using EcoRI and BglII diagnostic digests. Successful clones were verified by sequencing (Genewiz/Azenta).

### Adhesion Assay

Rotational adhesion assay was performed as described in (Artemenko et al., 2012). Briefly, 3x10^5^ cells (Figure 1) or 6x10^5^ cells (Figure 2) were plated in six-well tissue culture plates in 3 mL HL-5 with the appropriate antibiotics and grown overnight. The next day, media was aspirated and replaced with 3 mL fresh HL-5, cells were allowed to recover for one hour, and plates were shaken on an orbital shaker (New Brunswick Scientific G-33 for Figure 1; Fisherbrand Multi-Platform Shaker, cat no. 88-861-021 for Figure 2) at 150 RPM for one hour. After shaking, 1ml of media was collected from each well to count the floating cells. Remaining media was aspirated, and adherent cells were collected in 1ml of DB. % adhesion was calculated as (number of adherent cells)/(number of adherent + floating cells) x100%.

**Figure 1.**
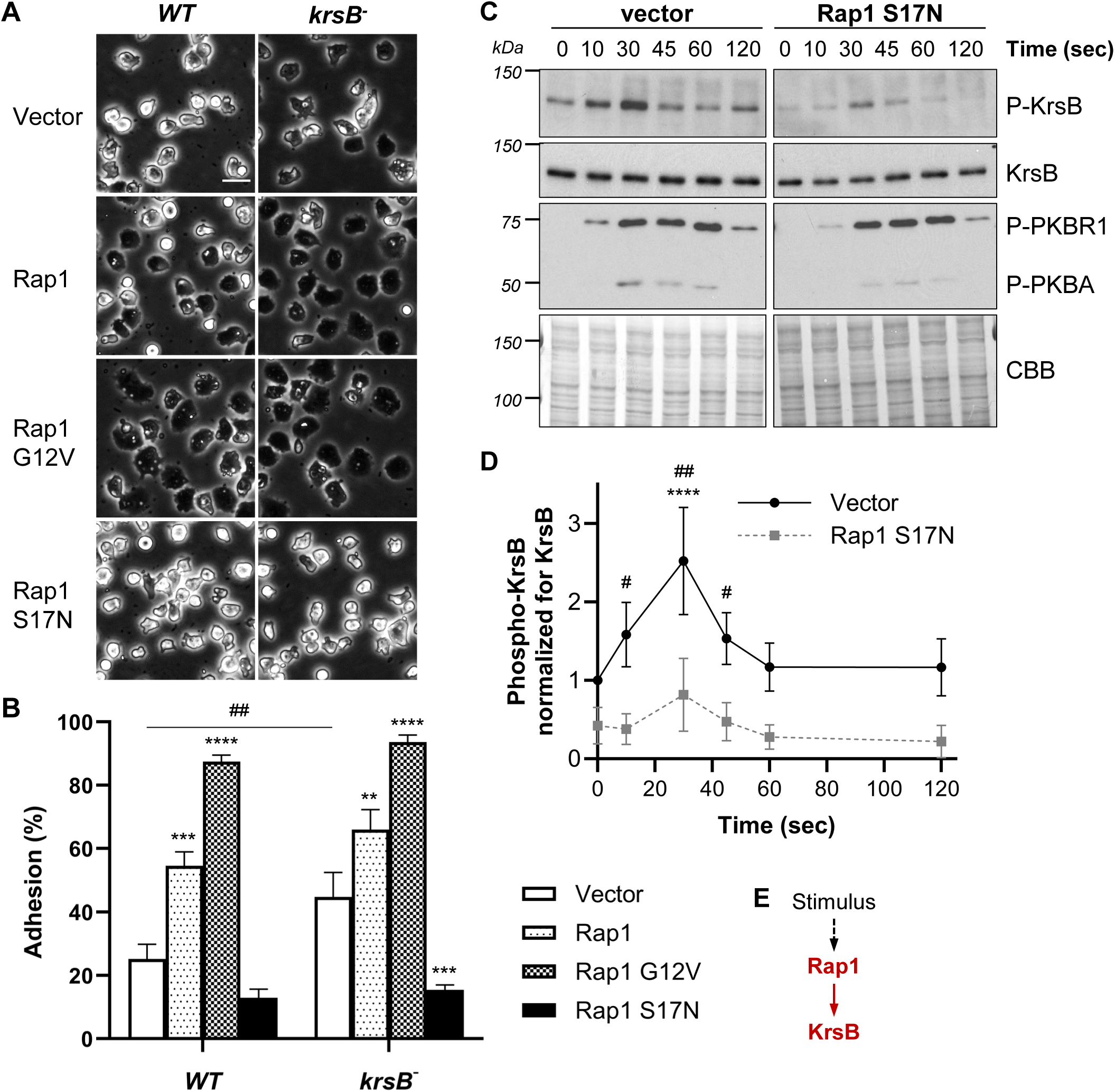
KrsB function in adhesion requires Rap1. (A-B) Vegetative wild-type (WT) or *krsB^−^* cells expressing Myc-tagged constitutively active (G12V), dominant negative (S17N), or wild-type Rap1 constructs were plated in HL-5 growth media and grown overnight. (A) Cells were imaged with phase microscopy at 200X magnification. Scale bar, 20 µm. (B) Media was replaced with fresh HL-5, cells were incubated for one hour and then subjected to a rotational adhesion assay at 150 RPM for 1 hr. % adhesion was calculated as the fraction of adherent over the total number of cell, times 100%. Data shown as mean ± SE of four independent experiments, each performed in duplicate. **P<0.01, ***P<0.001, ****P<0.0001 compared to vector control for a given cell line; ## P<0.01 compared to WT with the same construct using two-way ANOVA with Dunnett’s multiple comparisons test.. (C-D) Aggregation-competent WT or *krsB^−^* cells expressing vector or Rap1 S17N were stimulated with 10 µM cAMP and equal numbers of cells were lysed at the indicated time points after stimulation. Proteins were separated by SDS-PAGE and immunoblotted with antibodies against phospho-Mst1/2 (recognizes phospho-KrsB), KrsB, or phospho-PKC Thr410 (recognizes phospho-PKBR1 and phospho-PKBA). The blot was stained with Coomassie Brilliant Blue (CBB) to show protein amounts. (C) Representative immunoblots are shown. (D) Phospho-KrsB band intensity was quantified using ImageJ and normalized for the KrsB signal for the corresponding samples. Data shown as mean ± SE of three independent experiments. ****P<0.001 compared to time 0; #P<0.05, ##P<0.01 compared to Rap1 S17N at the same time point using two-way ANOVA with Dunnett’s multiple comparisons test.. (E) A cartoon highlighting Rap1-mediated activation of KrsB following chemoattractant stimulation.

**Figure 2.**
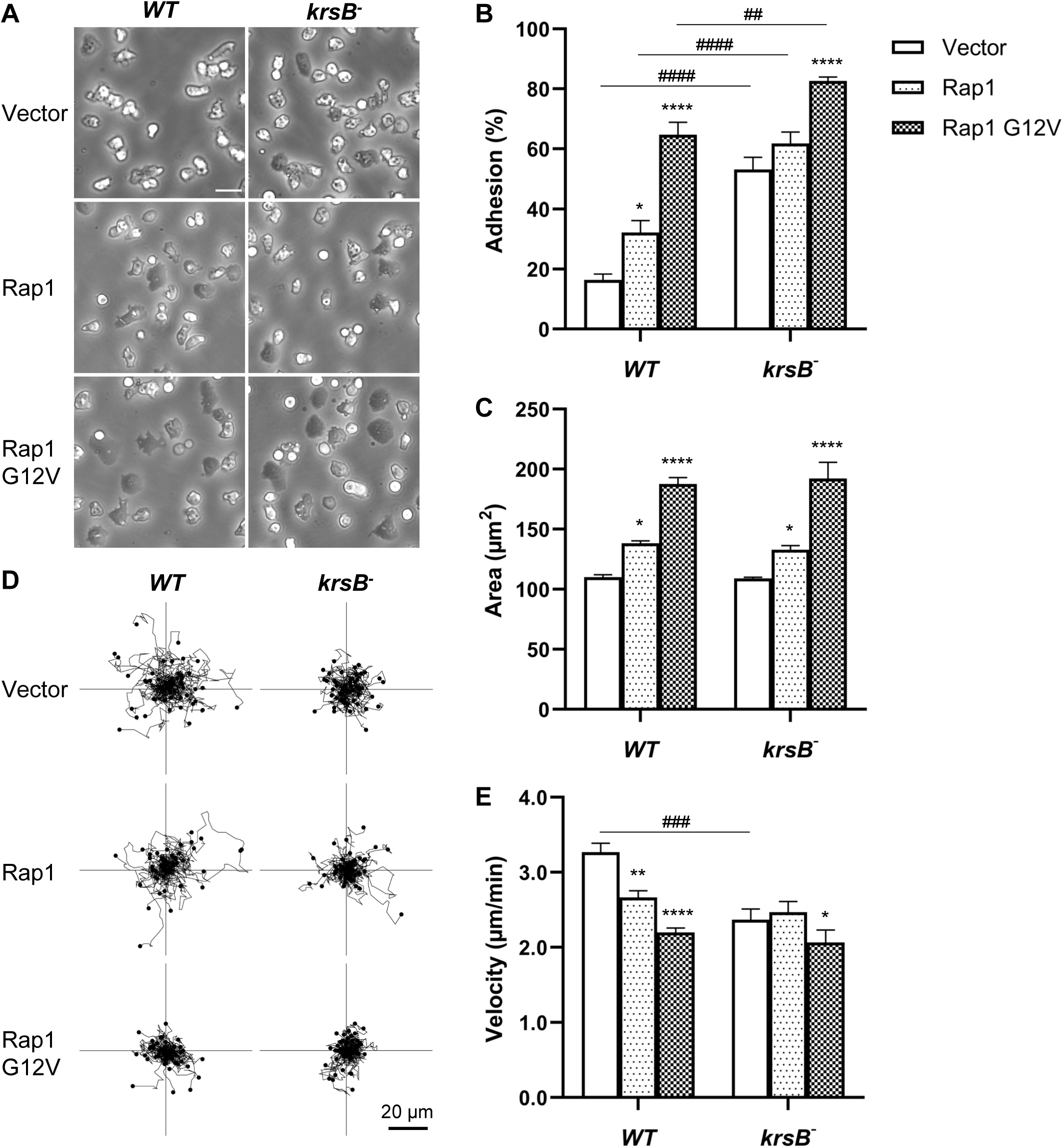
Rap1 does not require KrsB to mediate adhesion. Wild-type (WT) or krsB-null *D. discoideum* cells expressing RFP-tagged Rap1, constitutively active Rap1 G12V or empty vector were grown overnight in HL-5 media in tissue-culture coated plates. Media was replaced with fresh HL-5 one hour prior to analysis. (A) Representative phase-contrast images of cells were obtained at 200X magnification. Scale bar, 20 µm. (B) Cells were subjected to a rotational adhesion assay at 150 RPM for 1 hr and % adhesion was calculated as the fraction of adherent over the total number of cell, times 100%. Data shown as mean ± SE of four independent experiments. (C) Area was measured by manually tracing cell outlines in phase-contrast images from (A) using Fiji software. Data shown as mean ± SE of four independent experiments with 20 cells traced per experiment. (D-E) Random cell migration was measured by imaging cells under brightfield illumination at 200X magnification every 30 seconds for 40 cycles. Individual cells were manually tracked using Tracking Tool software. Representative tracks from one experiment are shown in (D). (E) Velocity data shown as mean ± SE of three independent experiments with 50 cells analyzed per experiment. *P<0.05, **P<0.01, ****P<0.0001 compared to vector control for a given cell line; ##P<0.01, ###P<0.001, ####P<0.0001 compared to WT with the same construct using two-way ANOVA with Dunnett’s multiple comparisons test.

### Migration Analysis and Morphological Assessment

For analysis of cells on tissue culture-coated plastic, 1x10^5^ cells were seeded in wells of a 24-well plate in 1 mL HL-5 with appropriate antibiotics and grown overnight. For analysis of cells on a glass surface, 5x10^4^ cells were seeded in wells of an 8-well LabTekII chambered coverglass in 500 µl HL-5 with appropriate antibiotics and grown overnight. The following day, cells were first imaged with phase-contrast illumination under a 20X objective using the Leica DMi1 microscope for morphological analysis. Individual cells were manually traced to determine cell area using Fiji ImageJ. The wells were then imaged simultaneously (up to 6 wells per experiment) with brightfield illumination under a 20X objective every 30 sec using Zeiss LSM700 Confocal Microscope. To improve the contrast of the images, the numerical aperture was lowered to 0.15. Czi files were converted to stacks of TIFF files using Fiji ImageJ and imported into TrackingTool Pro, where individual cells were tracked to determine the average velocity of the cells. A conversion ratio of 0.512 μm/pixel was used to calculate velocity in μm/min .

### Stimulation of Aggregation-Competent Cells

Aggregation-competent cells were prepared according to standard procedure (Artemenko et al., 2011). Briefly, 8x10^7^ cells were starved in 4 mL DB for one hour, followed by pulsing with 50 nM cAMP for 4 hours while rotating at 110 RPM on an orbital shaker. The success of differentiation was verified by checking levels of cAMP receptor cAR1 (see section on Immunoblotting), as well as by checking cell morphology and behavior under a microscope. For cAR1 lysates, 2x10^6^ cells were collected, resuspended in 40 µl 1X sample buffer (62.5 mM Tris·HCl pH 6.8, 2% SDS, 10% glycerol, 42 mM DTT, 0.01% bromophenol blue), and analyzed by immunoblotting (without boiling). To check cell morphology and behavior, 7x10^5^ cells were added to 1 mL of DB in a 24-well tissue culture coated plate at the end of the differentiation protocol, thoroughly pipetted up and down to disaggregate any clumps and observed after ∼20 min. Well-differentiated cells appeared elongated and began streaming.

Aggregation-competent cells used for protein localization analysis were seeded in chambers and analyzed as described below. For stimulation followed by cell lysis, cells were first basalated by incubation with 5 mM caffeine at 200 RPM for 30 min. After basalation, cells were washed with DB and resuspended at a final density of 4 x 10^7^ cells/ml. For time zero, 100 µl cells (4x10^6^ cells) were collected, mixed with 50 µl of 3X sample buffer and placed in a heating block at 98°C for 10 minutes. 1 mL of cells was mixed with 10 µl of 100 µM cAMP on an orbital shaker at 180 RPM. Samples were collected as above at 10, 30, 45, 60, and 120 seconds.

### Localization Analysis

For analysis of fluorescently tagged protein localization in vegetative cells, 5x10^4^ cells were seeded in wells of an 8-well LabTekII chambered coverglass in 500 ul HL-5 with appropriate antibiotics and grown overnight. The following day, cells were switched to 450 µl DB and incubated for one hour prior to imaging. Cells were imaged under 630X magnification, with either 488 nm or 555 nm laser with a Zeiss LSM700 confocal microscope. Specific settings for the laser intensity, pinhole, scan speed, number of scans averaged per image, and gain were kept constant between cell lines for a given experiment. Images were acquired every 3 seconds for 20 frames. After frame 5 cells (time 0) cells were stimulated by addition of 50 µl of 1 mM folic acid (100 µM final concentration). For analysis of fluorescently tagged protein localization in aggregation-competent cells, 10 µl of cells 5 hrs post-starvation (equivalent to 10^5^ cells) were seeded in 8-well LabTekII chambered coverglass with 440 µl DB and allowed to settle. Cells were imaged under 400X magnification, with a 488 nm laser with the UltraView spinning disk confocal microscope (DM 16000; Perkin-Elmer). Images were acquired every 5 seconds, and cells were stimulated by addition of 50 µl of 10 µM cAMP (1 µM final concentration) after frame 3 (time 0).

Response of fluorescently tagged proteins to chemoattractant stimulation was measured as the relative increase of fluorescence on the cortex/membrane of the cell. This was quantified by measuring the mean intensity of a 10x10 pixel square in the cytosol at every frame in Fiji ImageJ, normalizing the data for the mean intensity of the cytosol at time 0, and then inverting the data to show changes in the membrane intensity.

Basal localization of fluorescently tagged proteins was quantified by measuring the intensity of the protein signal on the membrane and expressing it as a ratio of the protein signal in the cytosol prior to stimulation (i.e., first frame of the timelapse). Membrane intensity was measured by outlining the cell along its perimeter, within 2-3 pixels from the edge, using the free tracing tool in Fiji ImageJ. The resulting area was converted to a line using the “Area to Line” function in the Selection tool (under Edit). Mean grey value of the line was then measured to determine the average pixel intensity along the perimeter, and was divided by the mean cytosolic intensity, which was measured as described above.

### Immunoblotting

Vegetative lysates for checking protein expression levels were obtained by lysing cells in 1X sample buffer at a final density of 2x10^7^ cells/mL and heating the samples at 98°C for 10 min. SDS-PAGE and immunoblotting were performed based on the standard procedures (Bio-Rad Laboratories, Inc., n.d.). Proteins were separated on a 7.5% resolving combined with 4% stacking polyacrylamide gels. For the KrsB phosphorylation analysis, proteins were resolved by a 4–15% Tris·HCl polyacrylamide gel (Criterion, Bio-Rad). Proteins were transferred onto a polyvinylidene fluoride membrane at 75V for 75 min. The membrane was blocked with 5% skim milk in TBST, followed by overnight incubation with a primary antibody in TBST with 5% BSA and 0.02% NaN_3_. The following primary antibodies were used: phosphoMst1/2 (Cell Signaling), phospho-PKCζ Thr410 (Cell Signaling), KrsB (Artemenko et al., 2012), Myc (Cell Signaling), cAR1 (Hereld et al., 1994), and mCherry (Invitrogen). Signal was detected by incubation with HRP-conjugated secondary antibody diluted blocking buffer and chemiluminescence detection using the Clarity^TM^ Western ECL Substrate kit (Bio-Rad).

### Statistical Analysis

All statistical analysis was performed in Prism (GraphPad). A two-way ANOVA with Dunnett’s multiple comparisons test was used to compare differences between multiple cell lines over time. P <0.05 was considered significant.

## Results

### KrsB function in adhesion requires Rap1

Since adhesion is a very robust and easily quantifiable phenotype for KrsB-null cells, we measured the effect of various Myc-tagged Rap1 constructs on adhesion of WT and KrsB-null cells (Figure 1A-B, S1). Consistent with published data, overexpression of Rap1 led to spreading and increased adhesion of WT cells by 2.3±0.3 fold (mean±SE; n=4). The increase in adhesion was even more pronounced in the presence of constitutively active Rap1 G12V (3.7±0.5 fold increase; mean ± SE; n=4). Also consistent with previous data, KrsB-null cells had significantly increased adhesion compared to WT cells (44±8 vs. 25±5%, respectively; mean±SE, n=4). Importantly, both Rap1 and Rap1 G12V further increased adhesion of krsB-null cells similarly to WT cells (1.6±0.3 and 2.3±0.4 fold increase, respectively; mean±SE; n=4), suggesting that Rap1 leads to changes in adhesion at least in part independently of KrsB. In contrast, expression of dominant negative Rap1 S17N led to reduced adhesion in WT cells by 2.0±0.2 fold (mean±SE, n=4). Importantly, it also reduced the increased adhesion of krsB-null cells to levels comparable to WT with Rap1 S17N (15±2% vs. 13±3%; mean±SE, n=4). Thus, the increased adhesion phenotype of KrsB-null cells appears to require functional Rap1.

If the KrsB-null phenotype depends on Rap1, it is possible that Rap1 somehow regulates KrsB activation. To address this possibility we examined KrsB phosphorylation following chemoattractant stimulation, which was previously shown to be indicative of KrsB activation (Artemenko et al., 2012). Stimulation of aggregation-competent wild-type cells with cAMP led to a transient increase in KrsB phosphorylation on Thr 176 with a peak response 30 sec after stimulation (Figure 1 C-D). Although the overall pattern of phosphorylation was similar in cells expressing Rap1 S17N, the extent of phosphorylation was much lower both basally (0.4±0.2 of the WT with vector signal at time 0) and following stimulation (2.5±0.7 vs. 2.0±0.1 fold increase at time 30 sec for vector vs. Rap1 S17N, respectively; mean±SE, n=3). Note that the reduced response was not due to reduced differentiation of cells expressing Rap1 S17N since levels of the cAMP receptor cAR1, which are used as an indicator of differentiation, were comparable between cells expressing empty vector and Rap1 S17N (Figure S2). Additionally, phosphorylation of another cAMP receptor target PKBR1/PKBA was also comparable between the two cell lines (Figure 1C). This suggests that Rap1 serves as an upstream activator of KrsB (Figure 1E).

### Rap1 regulates adhesion in the absence of KrsB

Initial findings demonstrated that Myc-tagged Rap1 can increase adhesion in the absence of KrsB (Figure 1A-B). To confirm our findings with Myc-tagged constructs, we generated Rap1 with an N-terminal RFP tag to allow visualization of Rap1. To make sure RFP-tagged Rap1 is functional, we tested the behavior of WT and KrsB-null cells expressing RFP-Rap1 or RFP-Rap1 G12V (Figure 2A). Since most of the cells were expressing the tagged constructs (Figure S3) we first performed population-level assays. Both Rap1 and Rap1 G12V significantly increased adhesion of WT cells (2.1±0.5 fold and 4.1±0.3, respectively; mean±SE, n=4; Figure 1B). In KrsB-null cells the effects were less pronounced with a 1.6±0.1 fold increase with Rap1 G12V expression, and no significant increase with Rap1. However, overall levels of adhesion achieved by KrsB-null cells with the Rap1 constructs were significantly higher than the WT counterparts.

The trends observed for adhesion were consistent with the overall morphological changes induced by Rap1 and Rap1 G12V. Both significantly increased the apparent area of cells observed using phase-contrast microscopy, although the relative increase was much greater for Rap1 G12V (Figure 2A, C). Similarly to adhesion, the change in area was comparable in WT and KrsB-null cells (1.71±0.05 and 1.8±0.1 fold increase for WT and KrsB-null with Rap1 G12V vs. vector, respectively; mean±SE, n=4), supporting the idea that Rap1 does not require KrsB for its function.

Increased spreading of cells expressing Rap1 and Rap1 G12V impaired the ability of WT cells to migrate, with the greatest reduction observed for Rap1 G12V (2.20±0.06 µm/min for Rap1 G12V vs. 3.3±0.1 µm/min for vector; Figure 2D-E). Surprisingly, despite a similar increase in spreading and adhesion of KrsB-null cells expressing Rap1 and Rap1 G12V, there was no significant reduction in velocity for these cells. This is likely because KrsB-null cells themselves were significantly slower than WT with a velocity of only 2.4±0.2 µm/min (mean±SE, n=3). The very low speeds seen for KrsB-null cells and WT cells with Rap1 G12V may reflect random fluctuations of the centroid during quantification instead of productive movement.

To improve the resolution of the migration assay, we analyzed the movement of cells on a glass surface, instead of tissue culture treated plastic (Figure 3). On this surface, Rap1 did not significantly increase the spreading of WT or KrsB-null cells, although Rap1 G12V still did so by 1.5 fold for both cell lines (Figure 3A-B). Under these less adhesive conditions, Rap1 did not significantly slow down WT cells, although Rap1 G12V did, consistent with the morphological assessment (3.0±0.1 for Rap1 G12V vs. 3.8±0.2 µm/min for vector; mean±SE, n=4; Figure 3C-D). In KrsB-null cells, Rap1 G12V did not significantly reduce the speed, possibly due to the low starting speed of KrsB-null cells as mentioned above. Surprisingly, despite lack of changes in the spreading due to Rap1 expression in KrsB-null cells, these cells moved significantly faster than KrsB-null cells expressing empty vector (3.6±0.2 vs. 2.70±0.08 µm/min; mean±SE, n=4). This was an unexpected finding suggesting that Rap1/KrsB interplay during migration may be more complex. Perhaps, Rap1 leads to more pseudopod production in the absence of KrsB, although this possibility remains to be explored.

**Figure 3.**
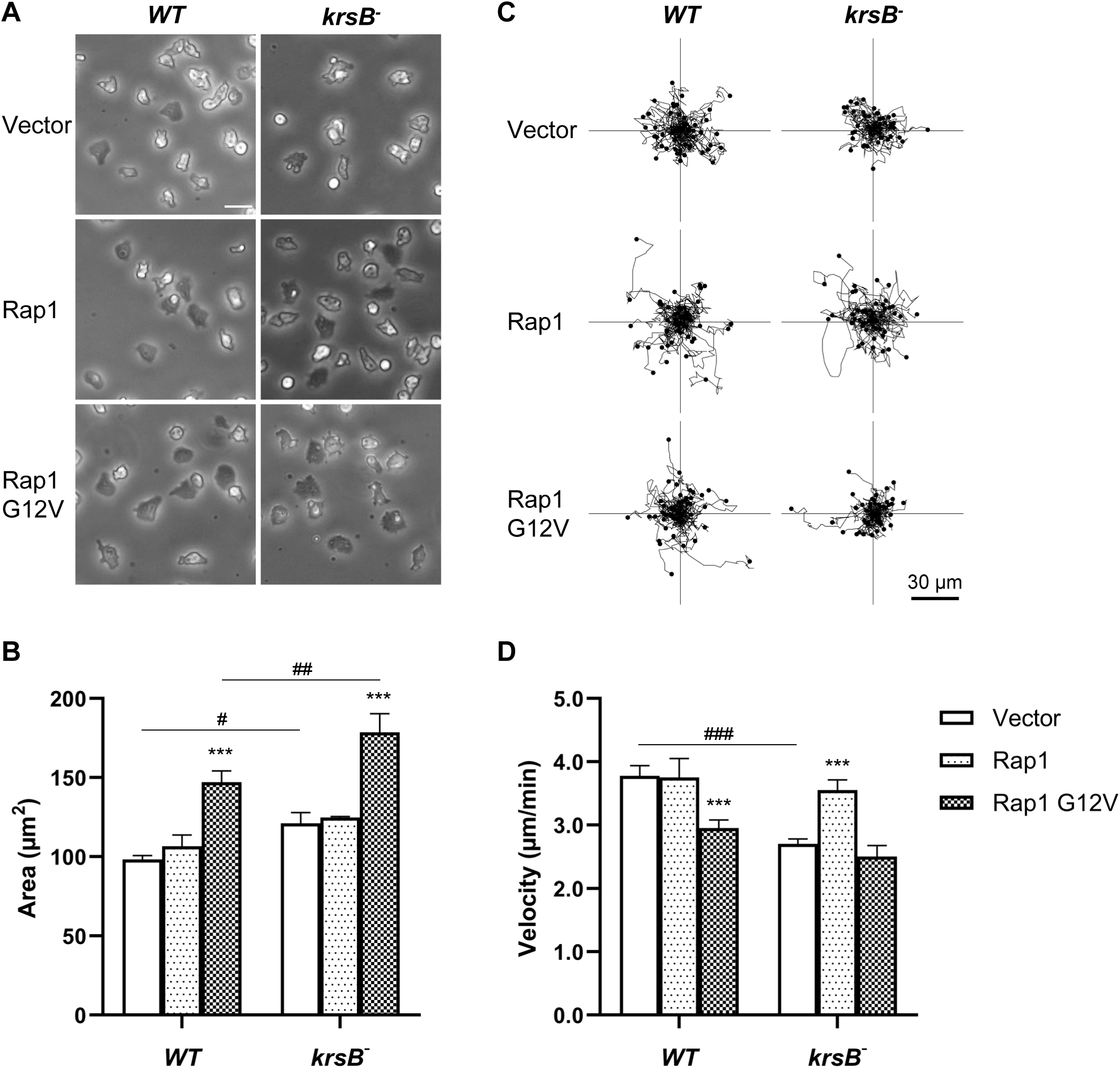
Rap1 does not require KrsB to mediate adhesion. Wild-type (WT) or krsB-null *D. discoideum* cells expressing RFP-tagged Rap1, constitutively active Rap1 G12V or empty vector were grown overnight in HL-5 media on a glass surface. (A) Representative phase-contrast images of cells were obtained at 200X magnification. Scale bar, 20 µm. (B) Area was measured by manually tracing cell outlines in phase-contrast images from (A) using Fiji software. Data shown as mean ± SE of three independent experiments with 30 cells traced per experiment. (C-D) Random cell migration was measured by imaging cells under brightfield illumination at 200X magnification every 30 seconds for 40 cycles. Individual cells were manually tracked using Tracking Tool software. Representative tracks from one experiment are shown in (C). (D) Velocity data shown as mean ± SE of four independent experiments with 50 cells analyzed per experiment. ***P<0.001 compared to vector control for a given cell line; #P<0.05, ##P<0.01, ###P<0.001 compared to WT with the same construct using two-way ANOVA with Dunnett’s multiple comparisons test.

### Rap1 activation is negatively regulated by KrsB

Since it appeared that Rap1 is able to affect adhesion in the absence of KrsB, but KrsB requires Rap1 for its function, we wanted to investigate if KrsB may be modulating Rap1 activity. We monitored Rap1 activation using GFP-labeled RalGDS, which preferentially binds to GTP-bound Rap1. This probe is known to transiently relocalize from the cytosol to the cortex following chemoattractant stimulation (Jeon et al., 2007). Consistently, we observed transient RalGDS accumulation on the membrane of WT cells following stimulation of vegetative cells with folic acid, with a peak response 9 sec after stimulation (Figure 4A-B). Although RalGDS relocalized in response to folic acid in KrsB-null cells as well, this response was weaker than in WT cells (1.19± 0.01 vs. 1.10±0.02 AU for WT vs. KrsB-null at 9 sec post-stimulation; mean±SE, n=53). Similar results were seen for aggregation-competent cells stimulated with cAMP (Figure S4), although we chose to focus our analysis on vegetative cells to avoid the confounding variable of differences in development between cell lines expressing various amounts of KrsB and Rap1.

**Figure 4.**
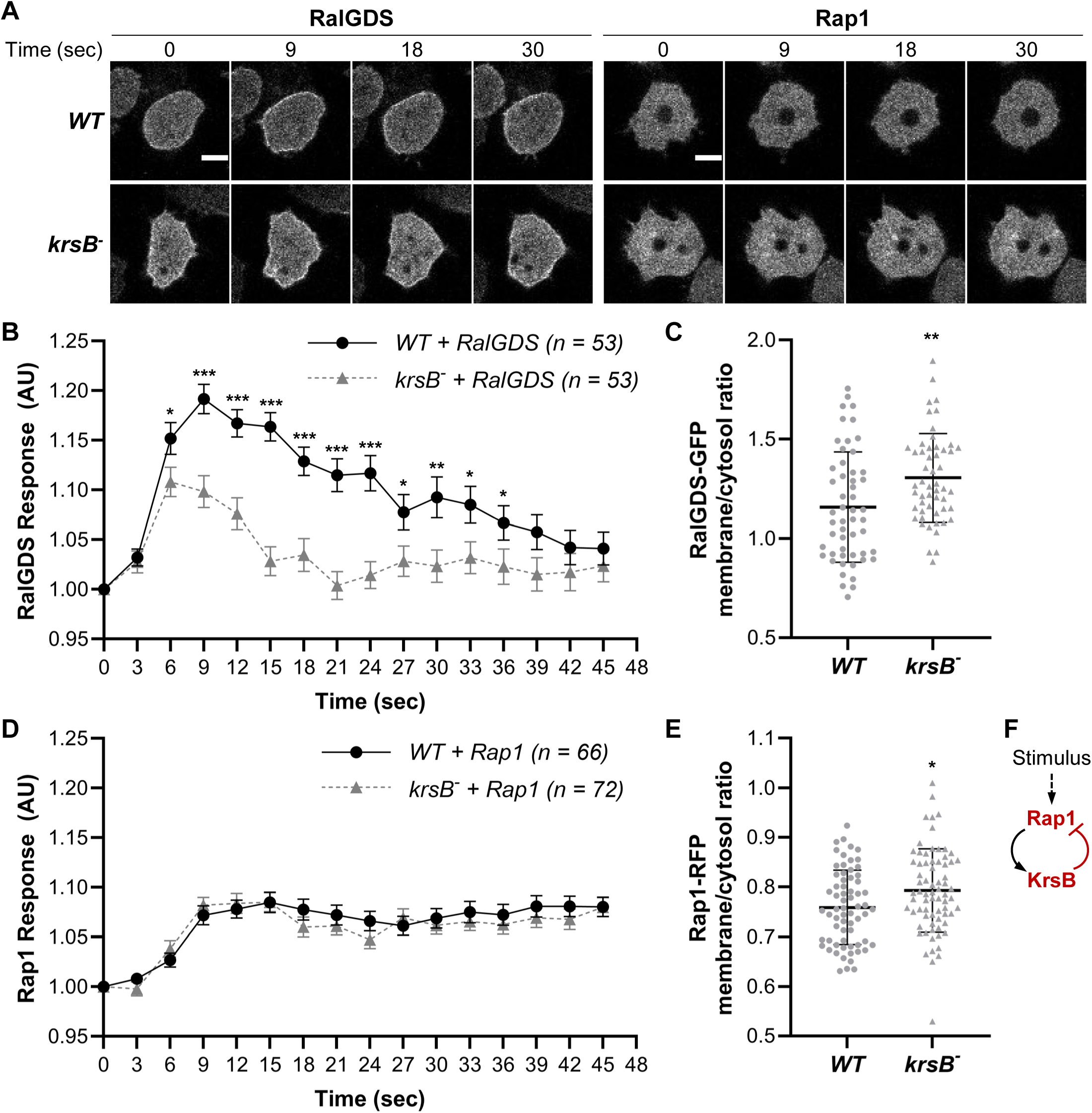
Comparison of Rap1 membrane localization in wild-type (WT) and KrsB-null cells basally and following chemoattractant stimulation. Vegetative *D. discoideum* cells expressing RalGDS-GFP or RFP-Rap1 were grown overnight in HL-5 media, switched to buffer, incubated for one hour, and imaged at 630X magnification every 3 sec using the 488 or 555 nm laser to detect the biosensor for active Rap1, RalGDS (A-C) or total Rap1 (A, D-E), respectively. After 5 frames, cells were stimulated with 100 µM folic acid (time 0). (A) Representative images are shown at the indicated times post-stimulation. Scale bar, 5 µm. (B, D) Membrane localization for RalGDS (B) or Rap1 (D) was quantified as the inverse of the mean intensity measured in a 10x10 pixel square in the cytosol at every frame using Fiji ImageJ software and normalized for the intensity at time 0. *P<0.05, **P<0.01, ***P<0.0001 compared to WT using two-way ANOVA with Dunnett’s multiple comparisons test. (C, D) Cells in images taken 12 sec prior to stimulation in (B) and (D) were manually traced along the edge of the cell (outermost 2-3 pixels), the area was converted to line and its intensity was measured using Fiji ImageJ software. This average membrane signal was normalized for the average cytosolic intensity determined as described in (B, D). All data shown as mean ± SE. Number of cells analyzed over three (B, C) or four (D, E) independent experiments is indicated in the key. *P<0.05, **P<0.01 compared to WT using an unpaired two-tailed t-test. (E) A cartoon highlighting a negative feedback loop between KrsB and Rap1 following chemoattractant stimulation.

Reduced RalGDS response in KrsB-null cells could suggest that either KrsB is required for Rap1 activation or that Rap1 activation is basally elevated, so that stimulation with a chemoattractant cannot further activate it. To distinguish between the two possibilities we quantified the ratio of membrane to cytosolic signal prior to stimulation. Consistent with the visual observations, membrane/cytosol levels of RalGDS were higher for KrsB-null cells compared to WT (1.31±0.03 vs. 1.16±0.04; mean±SE, n=53; Figure 4C), suggesting that Rap1 is basally activated to a greater extent in the absence of KrsB. Interestingly, basal levels of Rap1 itself (analyzed using RFP-Rap1) were also slightly, but significantly, elevated in the absence of KrsB (0.79±0.01 vs. 0.759±0.009; mean±SE, n=72 for KrsB-null and 66 for WT). Rap1showed some relocalization from the cytosol to the membrane following chemoattractant stimulation, but this response was not as pronounced compared to RalGDS and was not significantly different for WT and krsB-null cells (1.07±0.01 and 1.082±0.008 for WT and KrsB-null, respectively; mean±SE, n=72 for KrsB-null and 66 for WT). Note that even though the transient folic acid-induced recruitment of Rap1 to the membrane was weak, it was still measurable, for both Rap1 and to a small extent Rap1 G12V, but was absent for free cytosolic RFP (Figure S5A,C). There were no differences in basal membrane/cytosol ratio for constitutively active Rap1 or free RFP expressed in WT compared to KrsB-null cells (Figure S5B,D). Overall, this suggests that Rap1 activation is negatively regulated by KrsB (Figure 4E).

### Activation of downstream targets of Rap1 is increased in the absence of KrsB

If KrsB is a negative regulator of Rap1 activity, then activity of downstream targets of Rap1 should also be affected by KrsB. We examined the behavior of TalinB (TalB) and Phg2 in cells with or without KrsB following stimulation with a chemoattractant. TalB-GFP was basally localized on membrane protrusions and macropinosomes and was further recruited to the membrane following stimulation with folic acid, with the response being more pronounced in KrsB-null compared to WT cells (1.14±0.03 vs. 1.08±0.03, respectively, at 9 sec post-stimulation; mean±SE, n=35; Figure 5A-B). Although we also checked TalA, we did not observe any changes in localization in response to folic acid for this talin homolog (Figure S6). Phg2-GFP did not show any responsiveness to folic acid in WT cells, but showed significant recruitment to the membrane in cells lacking KrsB (Figure 5C-D). Unlike the transient response of RalGDS or TalB, membrane accumulation of Phg2-GFP remained high in cells lacking KrsB. This was consistent with the apparent increase in the basal levels of Phg2 on the membrane of KrsB-null cells, further suggesting that KrsB negatively regulates this Rap1 effector.

**Figure 5.**
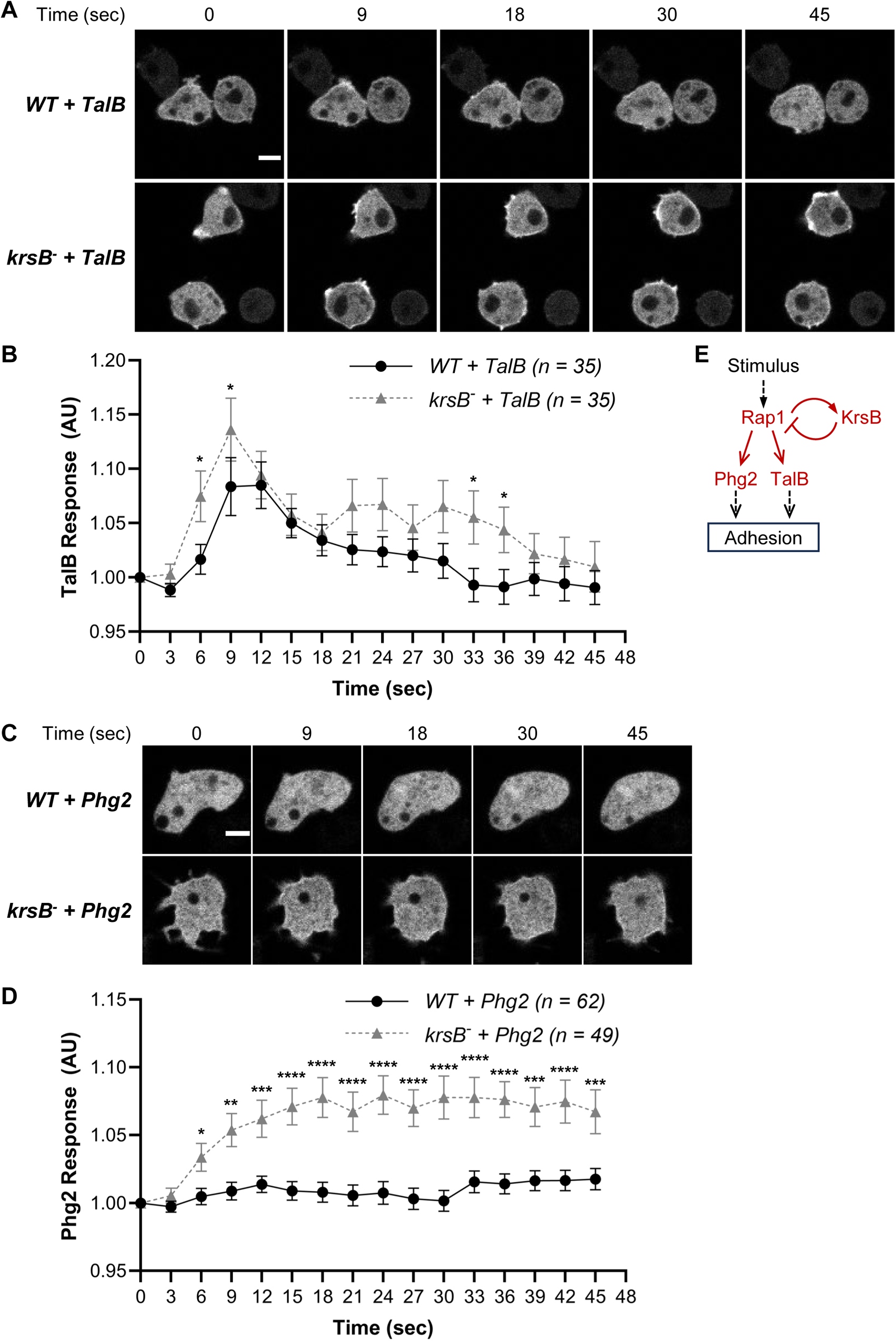
Comparison of localization of Rap1 effectors TalB and Phg2 in wild-type (WT) and KrsB-null cells following chemoattractant stimulation. Vegetative *D. discoideum* cells expressing TalB-GFP or Phg2-GFP were grown overnight in HL-5 media, switched to buffer, incubated for one hour, and imaged at 630X magnification every 3 sec using the 488 nm laser. After 5 frames, cells were stimulated with 100 µM folic acid (time 0). (A, C) Representative images are shown at the indicated times post-stimulation. Scale bar, 5 µm. (B, D) Membrane localization for TalB (B) or Phg2 (D) was quantified as the inverse of the mean intensity measured in a 10x10 pixel square in the cytosol at every frame using Fiji ImageJ software and normalized for the intensity at time 0. *P<0.05, **P<0.01, ***P<0.0001 compared to WT using two-way ANOVA with Dunnett’s multiple comparisons test. (E) Proposed model for the negative feedback loop limiting Rap1-mediated adhesion

## Discussion

Taken together, our data indicate that KrsB regulates adhesion by modulating the Rap1 pathway. We propose the following model: Rap1 activation, either due to random fluctuations of activity during random migration or downstream of chemoattractant stimulation, leads to elevated adhesion via Talin proteins and Phg2, which regulates contractility via myosin II. Rap1 also leads to activation of KrsB, which leads to the shut-off of Rap1-induced adhesion by reducing levels of Rap1 at the membrane. Thus, KrsB appears to be part of a negative feedback loop that limits the activity of the Rap1 pathway (Figure 5E).

Our data is consistent with Rap1 not requiring KrsB for its function, even though in some instances it appeared as if Rap1 was not as effective in KrsB-null cells compared to WT (e.g., adhesion data in Figure 2B). Based on the model above, KrsB-null cells likely have elevated endogenous Rap1 activity, which would explain why overexpression of Rap1 or Rap1 G12V does not change the phenotype of KrsB-null cells as much as in WT cells.

Our study did not address how Rap1 leads to increased KrsB activation. Although direct interaction between active Rap1 and KrsB was detected by mass spectrometry (Plak et al., 2016), this was not further investigated in this study. In mammalian cells, Mst1 activity and localization is regulated by a Rap1 effector, RAPL, and this activation is required for T lymphocyte polarization and integrin (LFA-1) clustering (Katagiri et al., 2006). There do not appear to be orthologs of RAPL in the *D. discoideum* genome, although a possibility of a functional ortholog cannot be ruled out. A notable difference is that Mst1 promotes adhesion of lymphocytes, whereas KrsB reduces adhesion of *D. discoideum.* It is possible that this is due to integrin-dependent vs. independent mode of adhesion in T lymphocytes vs. *D. discoideum*. It would be interesting to examine Mst1 function in integrin-independent migration of mammalian cells.

Alternatively, the differences between Mst1 and KrsB could be due to the unique structural features of KrsB, which has multiple regions homologous to domain III of Calpain at its C-terminus, or the absence of the SARAH domain, which is present in Mst1/2 but not KrsB (Artemenko et al., 2012).

It also remains to be seen whether KrsB activation occurs solely downstream of Rap1. We observed residual phosphorylation of KrsB in the presence of Rap1 S17N. This was not surprising even if Rap1 is essential for KrsB activation since lysates are obtained from a population of cells that likely includes ones that lack Rap1 S17N expression. Furthermore, even though Rap1 S17N acts in a dominant negative manner, some endogenous Rap1 function is likely preserved.

We demonstrated that KrsB alters Rap1 activation by affecting its localization. However, it remains unclear how KrsB blocks Rap1 from accessing the membrane and whether this effect is direct or indirect. In mammalian cells, Rap1 can be negatively regulated by phosphorylation by cAMP-dependent protein kinase-A (PKA) (Takahashi et al., 2013). Phosphorylation at the carboxyl terminus of Rap1 neutralizes the positive charges within the Rap1, which destabilizes the membrane localization and leads to sequestration of Rap1 by 14-3-3 in the cytosol (Takahashi et al., 2013). Since KrsB is a kinase, it is possible that it may directly phosphorylate Rap1 to cause its dissociation from the membrane, but this possibility remains to be examined. At this point we also cannot rule out that KrsB affects Rap1 indirectly, by modulating the activity of upstream GEF or GAPs.

Based on our study, negative regulation of Rap1 by KrsB is not selective for downstream targets of Rap1, since activation of both TalB and Phg2, which are not thought to be in a linear pathway (Hilbi and Kortholt, 2019), was affected. It might seem counterintuitive that activation of TalB and Phg2 is higher in KrsB-null cells considering that folic acid-stimulation of Rap1 (shown by RalGDS) is reduced in these cells. However, since the basal levels of Rap1 and active

Rap1 are higher in KrsB-null cells, even a relatively small increase in the active form of Rap1 on the membrane would be amplified, resulting in elevated activity levels of downstream effectors. Phg2 levels appeared to be higher at the cortex of KrsB-null cells basally as well. It should be noted that Phg2 has been previously shown to translocate to the membrane following cAMP stimulation (Jeon et al., 2007); however, we did not observe Phg2 translocation in response to folic acid stimulation in WT cells. This is likely because folic acid generally induces relatively weak responses in vegetative cells compared to the responses induced by cAMP in aggregation-competent cells. Observing a robust Phg2 response in KrsB-null cells, especially given this was in vegetative cells, is strong support that Phg2 activation is higher in the absence of KrsB.

It is unclear whether KrsB-mediated Rap1 regulation is only temporal or if there is a spatial component as well. Although we see overactivation of the Rap1 pathway all around the cell perimeter in cells lacking KrsB, it is likely that the inhibition of Rap1 would be restricted to certain regions of the cells, similarly to where Rap1 itself is activated (leading edge, protrusions). Although our previous study did not see any specific localization of KrsB in the cell (Artemenko et al., 2012), if Rap1 is responsible for activation of KrsB, it would make sense that active KrsB would follow the same pattern as active Rap1.

Another remaining question is whether KrsB has functions outside of the Rap1 pathway. Since Rap1 S17N completely rescued increased adhesion of KrsB-null cells, it seems that whichever way KrsB modulates adhesion, it happens upstream of the S17N effect, so that in the absence of Rap1, KrsB can no longer function to modulate adhesion. It is interesting to note that cells lacking KrsB and expressing empty vector migrated as poorly as WT cells expressing Rap1 G12V even though these cells were not as spread or as adhesive. This could suggest that cells lacking KrsB may have impaired function of additional pathways beyond Rap1. However, it is also possible that there are differences between overactivation of the endogenous Rap1 pathway in KrsB-null cells compared to the overexpression of constitutively active Rap1 G12V. Some evidence for this can be found in the migration behavior of KrsB-null cells expressing Rap1, which migrated faster on a glass surface compared to control despite having the same area. Further studies are needed to parse out the precise relationship between spreading and migration in these cells.

Overall, our study demonstrates that Rap1 activation leads to activation of its own shut-down mechanism via KrsB, although questions remain on the molecular mechanism of both KrsB phosphorylation downstream of Rap1 and Rap1 sequestration away from the membrane downstream of KrsB. The negative feedback loop between Rap1 and KrsB may provide dynamic regulation of cell-substrate adhesion that is necessary for rapid amoeboid-type migration.

## Acknowledgements

The authors would like to thank Dr. Arjan Kortholt for helpful discussions and critical reading of the manuscript, as well as Dr. Richard Firtel, Dr. Arjan Kortholt, and the Dicty Stock Center for providing expression constructs and strains used in this study. This work was supported by NIH grant R35 GM118177 (to P.N.D.) and NSF-RUI grant no. 1817378 (to Y.A.).

## Notes

### Competing Interest Statement

The authors have declared no competing interest.

## References

Artemenko, Y., Batsios, P., Borleis, J., Gagnon, Z., Lee, J., Rohlfs, M., et al. (2012). Tumor suppressor Hippo/MST1 kinase mediates chemotaxis by regulating spreading and adhesion. Proc. Natl. Acad. Sci. U.S.A. 109, 13632–13637. doi: 10.1073/pnas.1211304109

Artemenko, Y., Lampert, T. J., and Devreotes, P. N. (2014). Moving towards a paradigm: common mechanisms of chemotactic signaling in Dictyostelium and mammalian leukocytes. Cell. Mol. Life Sci. 71, 3711–3747. doi: 10.1007/s00018-014-1638-8

Artemenko, Y., Swaney, K. F., and Devreotes, P. N. (2011). Assessment of development and chemotaxis in Dictyostelium discoideum mutants. Methods Mol. Biol. 769, 287–309. doi: 10.1007/978-1-61779-207-6_20

Bio-Rad Laboratories, Inc. (n.d.). Bulletin 6040 Ver C: A Guide to Polyacrylamide Gel Electrophoresis and Detection. Available at: https://www.bio-rad.com/webroot/web/pdf/lsr/literature/Bulletin_6040.pdf

Bozzaro, S. (2019). The past, present and future of Dictyostelium as a model system. Int. J. Dev. Biol. 63, 321–331. doi: 10.1387/ijdb.190128sb

Bukharova, T., Weijer, G., Bosgraaf, L., Dormann, D., Van Haastert, P. J., and Weijer, C. J. (2005). Paxillin is required for cell-substrate adhesion, cell sorting and slug migration during *Dictyostelium* development. J. Cell Sci. 118, 4295–4310. doi: 10.1242/jcs.02557

Cornillon, S., Froquet, R., and Cosson, P. (2008). Involvement of Sib Proteins in the Regulation of Cellular Adhesion in *Dictyostelium discoideum*. Eukaryot. Cell 7, 1600–1605. doi: 10.1128/EC.00155-08

Gaudet, P., Pilcher, K. E., Fey, P., and Chisholm, R. L. (2007). Transformation of Dictyostelium discoideum with plasmid DNA. Nat. Protoc. 2, 1317–1324. doi: 10.1038/nprot.2007.179

Hereld, D., Vaughan, R., Kim, J. Y., Borleis, J., and Devreotes, P. (1994). Localization of ligand-induced phosphorylation sites to serine clusters in the C-terminal domain of the Dictyostelium cAMP receptor, cAR1. J. Biol. Chem. 269, 7036–7044.

Hibi, M., Nagasaki, A., Takahashi, M., Yamagishi, A., and Uyeda, T. Q. P. (2004). Dictyostelium Discoideum Talin A is Crucial for Myosin II-Independent and Adhesion-Dependent Cytokinesis. J. Muscle. Res. Cell Motil. 25, 127–140. doi: 10.1023/B:JURE.0000035842.71415.f3

Hilbi, H., and Kortholt, A. (2019). Role of the small GTPase Rap1 in signal transduction, cell dynamics and bacterial infection. Small GTPases 10, 336–342. doi: 10.1080/21541248.2017.1331721

Jeon, T. J., Lee, D.-J., Merlot, S., Weeks, G., and Firtel, R. A. (2007). Rap1 controls cell adhesion and cell motility through the regulation of myosin II. J. Cell Biol. 176, 1021– 1033. doi: 10.1083/jcb.200607072

Katagiri, K., Imamura, M., and Kinashi, T. (2006). Spatiotemporal regulation of the kinase Mst1 by binding protein RAPL is critical for lymphocyte polarity and adhesion. Nat. Immunol. 7, 919–928. doi: 10.1038/ni1374

Lämmermann, T., Bader, B. L., Monkley, S. J., Worbs, T., Wedlich-Söldner, R., Hirsch, K., et al. (2008). Rapid leukocyte migration by integrin-independent flowing and squeezing. Nature 453, 51–55. doi: 10.1038/nature06887

Loomis, W. F., Fuller, D., Gutierrez, E., Groisman, A., and Rappel, W.-J. (2012). Innate Non-Specific Cell Substratum Adhesion. PLoS ONE 7, e42033. doi: 10.1371/journal.pone.0042033

Mijanović, L., and Weber, I. (2022). Adhesion of Dictyostelium Amoebae to Surfaces: A Brief History of Attachments. Front. Cell Dev. Biol. 10, 910736. doi: 10.3389/fcell.2022.910736

Parlani, M., Jorgez, C., and Friedl, P. (2023). Plasticity of cancer invasion and energy metabolism. Trends Cell Biol. 33, 388–402. doi: 10.1016/j.tcb.2022.09.009

Plak, K., Pots, H., Van Haastert, P. J. M., and Kortholt, A. (2016). Direct Interaction between TalinB and Rap1 is necessary for adhesion of Dictyostelium cells. BMC Cell Biol. 17, 1. doi: 10.1186/s12860-015-0078-0

Pourjafar, M., and Tiwari, V. K. (2024). Plasticity in cell migration modes across development, physiology, and disease. Front. Cell Dev. Biol. 12, 1363361. doi: 10.3389/fcell.2024.1363361

Smith, A., Carrasco, Y. R., Stanley, P., Kieffer, N., Batista, F. D., and Hogg, N. (2005). A talin-dependent LFA-1 focal zone is formed by rapidly migrating T lymphocytes. J. Cell Biol. 170, 141–151. doi: 10.1083/jcb.200412032

Takahashi, M., Dillon, T. J., Liu, C., Kariya, Y., Wang, Z., and Stork, P. J. S. (2013). Protein Kinase A-dependent Phosphorylation of Rap1 Regulates Its Membrane Localization and Cell Migration. J. Biol. Chem. 288, 27712–27723. doi: 10.1074/jbc.M113.466904

Tsujioka, M., Yoshida, K., Nagasaki, A., Yonemura, S., Müller-Taubenberger, A., and Uyeda, T. Q. P. (2008). Overlapping Functions of the Two Talin Homologues in *Dictyostelium*. Eukaryot. Cell 7, 906–916. doi: 10.1128/EC.00464-07

