## Supplemental Figures for "A negative feedback loop between small GTPase Rap1 and mammalian tumor suppressor homolog KrsB regulates cell-substrate adhesion in *Dictyostelium*"

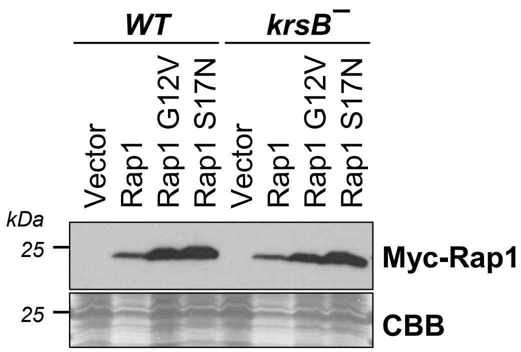


**Figure S1. Expression levels of Myc-tagged Rap1 constructs.** Equal numbers of vegetative wild-type (WT) or *krsB*¯ cells expressing Myc-tagged constitutively active (G12V), dominant negative (S17N), or wild-type Rap1 constructs were lysed, proteins were separated by SDS-PAGE and immunoblotted with antibodies against Myc. The blot was stained with Coomassie Brilliant Blue (CBB) to show protein amounts. A representative immunoblot is shown. This figure corresponds to Figure 1A-B.

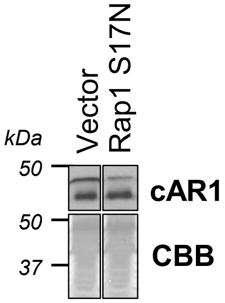


**Figure S2. Dominant negative Rap1 (S17N) expression does not disrupt development of wild-type (WT) cells.** Equal numbers of aggregation-competent WT cells expressing Myc-tagged Rap1 S17N or empty vector were lysed, proteins were separated by SDS-PAGE and immunoblotted with antibodies against cAMP receptor cAR1. The blot was stained with Coomassie Brilliant Blue (CBB) to show protein amounts. A representative immunoblot is shown. This figure corresponds to Figure 1C-D.

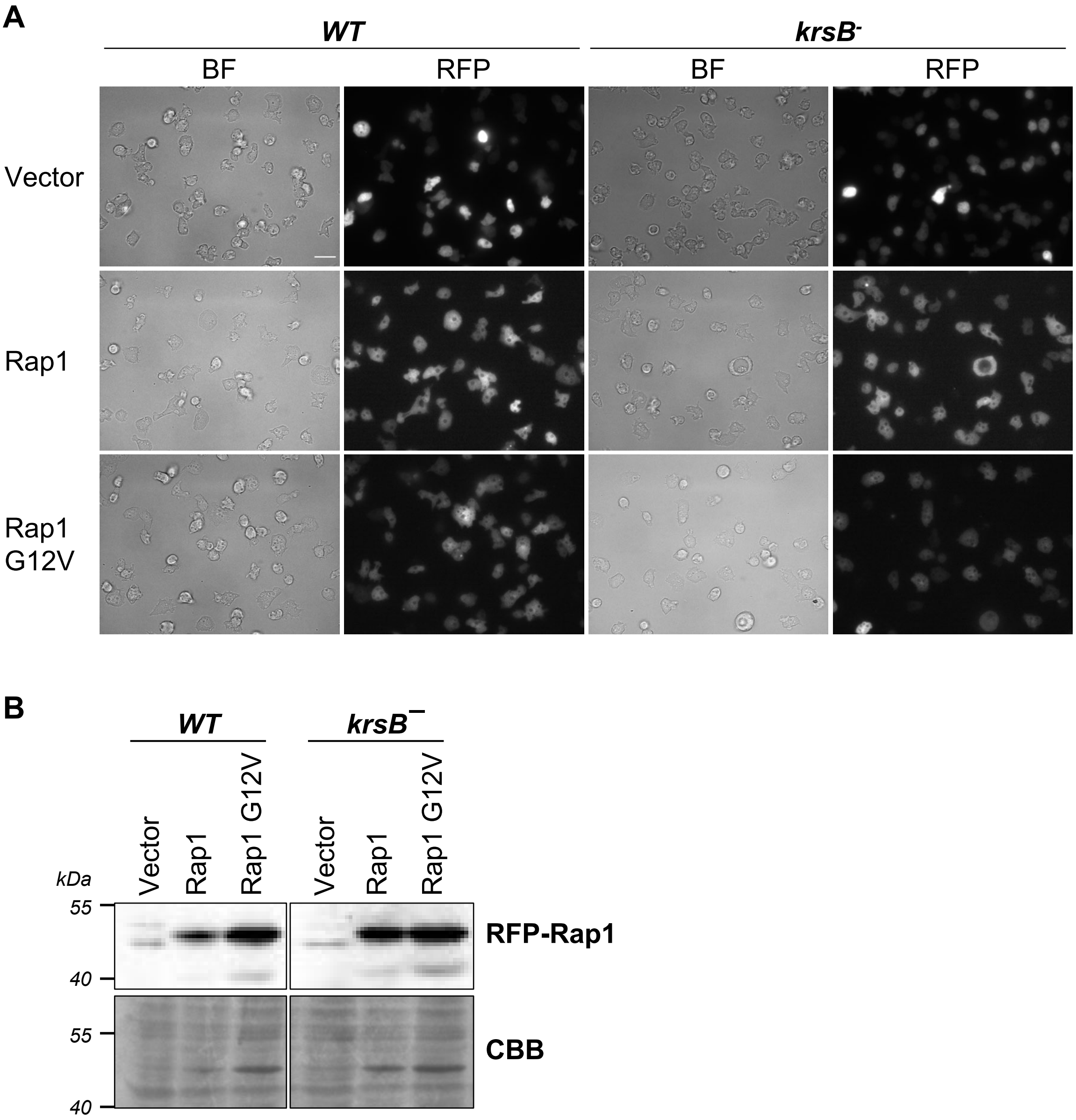


**Figure S3. Expression levels of RFP-tagged Rap1 constructs.** Wild-type (WT) or krsB-null *D. discoideum* cells were transformed with RFP-tagged constitutively active (G12V) or wild-type Rap1 constructs or empty vector encoding RFP. (A) Cells were plated glass-bottom chambers in buffer, incubated for one hour, and imaged with brightfield illumination or epifluorescence with an RFP filter set at 630X magnification. An entire field is shown. Scale bar, 20 µm. (B) Equal numbers of cells were lysed, proteins were separated by SDS-PAGE and immunoblotted with antibodies against mCherry. The blot was stained with Coomassie Brilliant Blue (CBB) to show protein amounts. A representative immunoblot is shown. This figure corresponds to Figures 2-3.

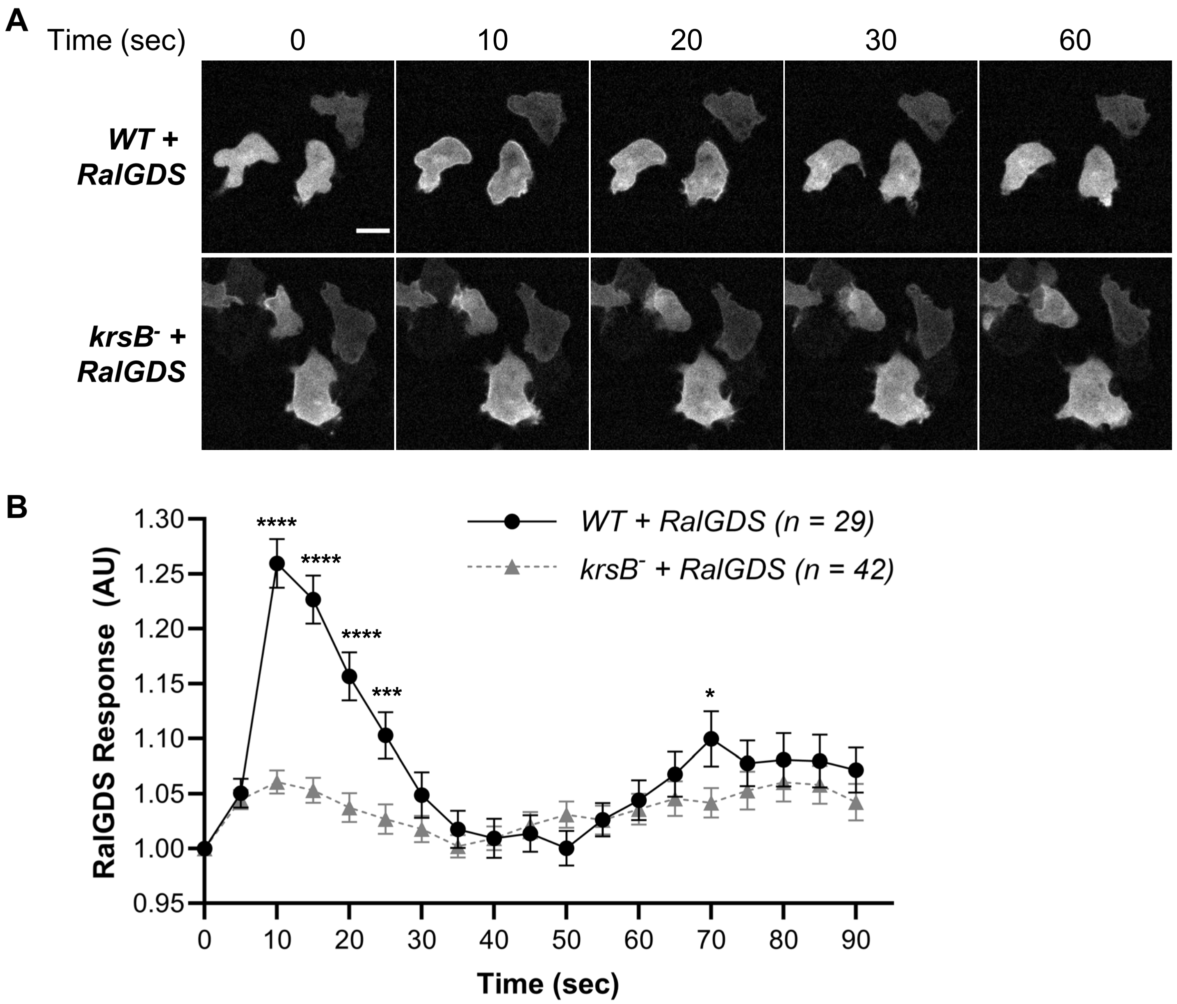


**Figure S4. Analysis of cAMP-induced Rap1 activation.** Aggregation-competent wild-type or *krsB*¯ cells expressing a biosensor for Rap1 activation (RalGDS-GFP) were imaged at 400X magnification every 5 sec using the 488 nm laser. After 5 frames, cells were stimulated with 1 µM cAMP. (A) Representative images are shown at the indicated times post-stimulation. Scale bar, 10 µm. (B) Membrane localization of RalGDS was quantified as the inverse of the mean intensity measured in a square in the cytosol at every frame using Fiji ImageJ software and normalized for the intensity at time 0. Data shown as mean ± SE. Number of cells analyzed is indicated in the key. *P<0.05, ***P<0.001, ***P<0.0001 compared to WT using two-way ANOVA with Dunnett's multiple comparisons test.

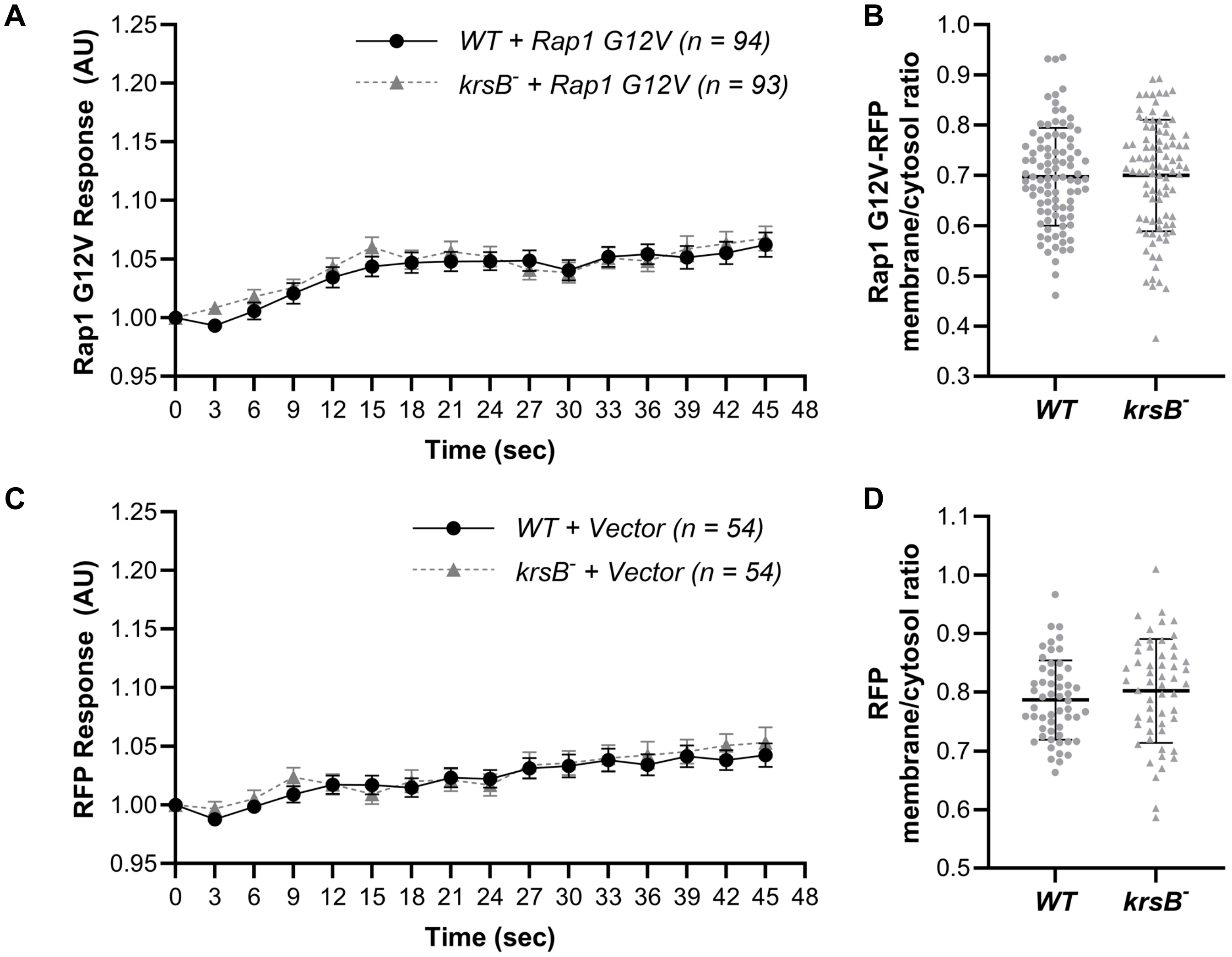


**Figure S5. Comparison of Rap1 G12V and free RFP membrane localization in wild-type (WT) and KrsB-null cells basally and following chemoattractant stimulation.** Vegetative *D. discoideum* cells expressing RFP-Rap1 G12V or vector encoding free RFP were grown overnight in HL-5 media, switched to buffer, incubated for one hour, and imaged at 630X magnification every 3 sec using the 555 nm laser. After 5 frames, cells were stimulated with 100 μM folic acid (time 0). (A, C) Membrane localization for Rap1 G12V (A) or RFP (C) was quantified as the inverse of the mean intensity measured in a 10x10 pixel square in the cytosol at every frame using Fiji ImageJ software and normalized for the intensity at time 0. Number of cells analyzed over five (A) or three (C) independent experiments is indicated in the key. (B, D) Cells in images taken 12 sec prior to stimulation in (A) and (C) were manually traced along the edge of the cell (outermost 2-3 pixels), the area was converted to line and its intensity was measured using Fiji ImageJ software. This average membrane signal was normalized for the average cytosolic intensity determined as described in (A, C). Data shown as mean ± SE. The numbers of cells analyzed were the same as in (A) and (C), except for *krsB*$^-$ cells expressing Rap1 G12V, where only 92 cells were analyzed for basal localization.

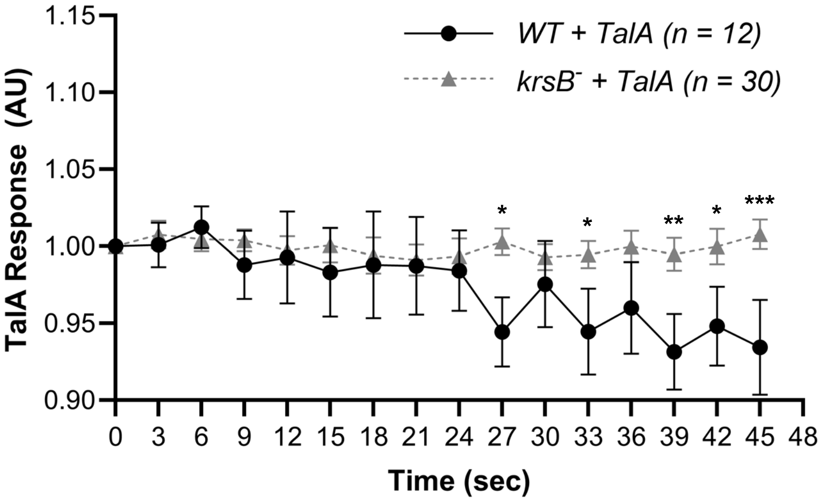


**Figure S6. Comparison of localization of Rap1 effector TalA in wild-type (WT) and KrsB-null cells following chemoattractant stimulation.** Vegetative *D. discoideum* cells expressing TalA-GFP were grown overnight in HL-5 media, switched to buffer, incubated for one hour, and imaged at 630X magnification every 3 sec for 20 cycles using the 488 nm laser. After 5 frames, cells were stimulated with 100 µM folic acid. Membrane localization for was quantified as the inverse of the mean intensity measured in a 10x10 pixel square in the cytosol at every frame using Fiji ImageJ software and normalized for the intensity at time 0. *P<0.05, **P<0.01, ***P<0.001 compared to WT using two-way ANOVA with Dunnett's multiple comparisons test.
